# Automatic sorting system for large calcium imaging data

**DOI:** 10.1101/215145

**Authors:** Takashi Takekawa, Hirotaka Asai, Noriaki Ohkawa, Masanori Nomoto, Reiko Okubo-Suzuki, Khaled Ghandour, Masaaki Sato, Yasunori Hayashi, Kaoru Inokuchi, Tomoki Fukai

## Abstract

It has become possible to observe neural activity in freely moving animals via calcium imaging using a microscope, which could not be observed previously. However, it remains difficult to extract the dynamics of nerve cells from the recorded imaging data. In this study, we greatly improved the stability, and robustness of the cell activity estimation method via non-negative matrix decomposition with shrinkage estimation of the baseline. In addition, by improving the initial state of the iterative algorithm using a newly proposed method to extract the shape of the cell via image processing, a solution could be obtained with a small number of iterations. These methods were applied to artificial and real data, and their effectiveness was confirmed.

## INTRODUCTION

Calcium imaging is used to visualize the activity of large neural populations, and it is becoming a standard technique for recording neural activity in normally behaving animals. Recent advances in genetic engineering and recording technologies have significantly improved the signal intensity of calcium-binding fluorescent indicators, increasing the spatial and temporal resolutions of imaging data (Chen, et al., 2013). Some calcium indicators are sufficiently sensitive to detect action potentials in vivo (Chen, et al., 2013). Furthermore, we can currently image neuronal activity in vivo in a wider region of the brain for longer periods (Cotton, et al., 2013; Ahrens, et al., 2013; Prevedel, et al., 2014; Diego-Andilla & Hamprecht, 2013). Accordingly, the demand for efficient and accurate methods for detecting the spatial locations and shapes (footprints) of individual cells together with their temporal firing patterns from imaging data has increased.

Conventionally, the spatial locations of individual cells are first identified in the imaging data. Then, the shape of each cell is extracted from these data and marked as the region of interest (ROI) or footprint, which usually occupies multiple pixels. Knowing that cells occupy localized spatial regions, we may use local correlations of neighboring pixels (Smith & Häusser, 2010) or principal component analysis (Mukamel, et al., 2009) to attempt an automatic extraction of their footprints. However, the accuracy of these methods is not always sufficient, and time-consuming manual extraction via visual inspection remains the primary modality for this purpose. At the second step, the spiking activity of each neuron is estimated from slowly varying image intensity, which is usually averaged over the imaged pixels belonging to each footprint to improve the signal-to-noise (S/N) ratio (Smith & Häusser, 2010). Several methods exist for deconvoluting spike sequences (Grewe, et al., 2010; Oñativia, et al., 2013; Theis, et al., 2016), among which Markov chain Monte Carlo methods (Pnevmatikakis, et al., 2013) often work accurately. However, the accuracy of these methods is significantly degraded when the footprints of different cells spatially overlap, and this often occurs because of the projection of the 3D neural circuit structure onto a 2D image space or a low spatial resolution in optical recordings. Separating spatially overlapping cells represents a challenge even with state-of-the-art methods for analyzing calcium imaging data.

To handle this problem, it has been proposed to represent a recorded image using the sum of the products of spatial (footprints) and temporal components (intensity changes). Independent component analysis was examined for this purpose (Mukamel, et al., 2009), but it was not sufficient for decorrelating signals from spatially overlapping cells. Some nonlinear methods based on multilevel sparse matrix factorization (Diego-Andilla & Hamprecht, 2013) or nonnegative matrix factorization (NMF) (Maruyama, et al., 2014) improved the decorrelation performance. However, these methods did not consider the dynamics of neuronal firing or the intracellular calcium concentration. As a result, these methods were impaired by the noise effect that deteriorates the accuracy in decomposing optical signals into activity-dependent and noise components.

In addition to the aforementioned problem, another technical challenge is finding an adequate estimate of the baseline level of imaging data. Only the relative intensity of optical signals from different cells is meaningful in calcium imaging recordings. However, because the baseline signal intensity of silent cells is unknown, it is difficult to unambiguously distinguish the faint intensity changes caused by actual biological processes, such as neuronal firing at low frequencies, from the stochastic fluctuations arising from the spatial inhomogeneity of recorded materials or temporal fluctuations during optical recordings.

Against this background, a framework called constrained nonnegative matrix factorization (CNMF), which introduced calcium dynamics and the sparsity of spikes into NMF, was recently proposed (Pnevmatikakis, et al., 2013; Pnevmatikakis, et al., 2016). In addition, enhanced CNMF, which considers baselines inadequate for data with significant overlap, has been described. Because CNMF fixes time and spatial elements and optimizes the remaining elements, it is important to set good initial conditions and ensure convergence via iteration. However, in the current situation, ad hoc processing is performed on the result with few iterations and corrected, and coverage is not guaranteed; thus, doubt remains regarding the validity and stability of the obtained result. As the data size further increases in the future, strategies for evaluating the data will become more important.

In this study, we propose a novel initialization method for footprints and two simple but powerful extensions for CNMF. The proposed initialization method using a Laplacian of Gaussian (LoG) filter can acquire footprints with small false-negative results even if the data contain cells with low S/N ratios and firing frequencies. The introduction of shrinkage estimation for baseline to CNMF improves the stability of the iterative steps of CNMF. Simultaneous estimation of the cell components and baseline also improves the accuracy and stability, especially when data contain strongly overlapped the cells. By combining the initialization and iterative methods, it is possible to accurately detect temporal and spatial structures for artificial and real data, even for cells with low firing rates of signals that were overlooked using conventional methods.

## RESULTS

### LoG filters can reliably detect low-firing-rate cells

Using the footprint as the initial condition, in most studies, cell candidates are selected on the basis of statistics over the frames. However, cells with low SN ratios and/or firing rates are difficult to detect using such a method. Therefore, we developed a method to detect cell candidates for each frame. The LoG filter, which is used for blob detection in the field of image processing, is utilized to emphasize a closed region of a specific size from an image with substantial noise, and it is possible to efficiently detect a cell candidate from each frame.

In this study, we evaluated the proposed method to initialize cell footprints using artificial data. The composition of the artificial data is shown in Fig. 1. In this example, the activities of 200 cells with strong overlap were simulated. Cell size and signal intensity also significantly fluctuated (see Simulation Data in Method Details). In the proposed method, LoG filters with multiple scales are first applied to each frame (i.e., Fig. 2A), as shown in Fig. 2B. The peaks in LoG-filtered images are detected within three dimensions of position and scale (Fig. 2C).

**Figure 1.**
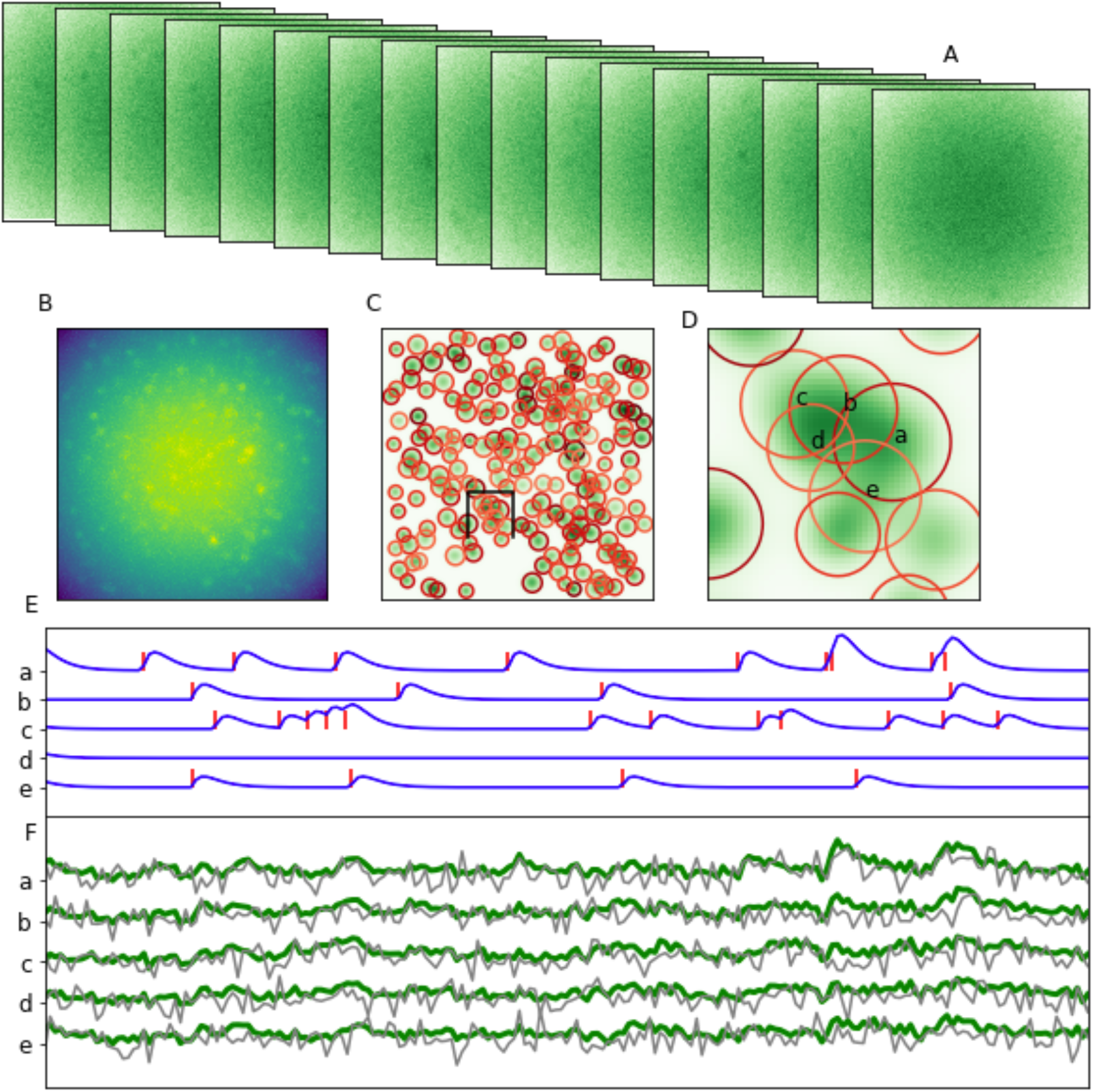
Explanation of the model concept using artificial data. Observed motion picture data are presented. The maximum value of imaging data was obtained in the time direction. Contour diagrams were overlaid onto cell shape data. Expanding a part of C. The spike activity and change in the calcium concentration in each cell are presented, in addition to the time series observed for each cell. Gray denotes the position of the peak. Green denotes the average value by region of interest.

**Figure 2.**
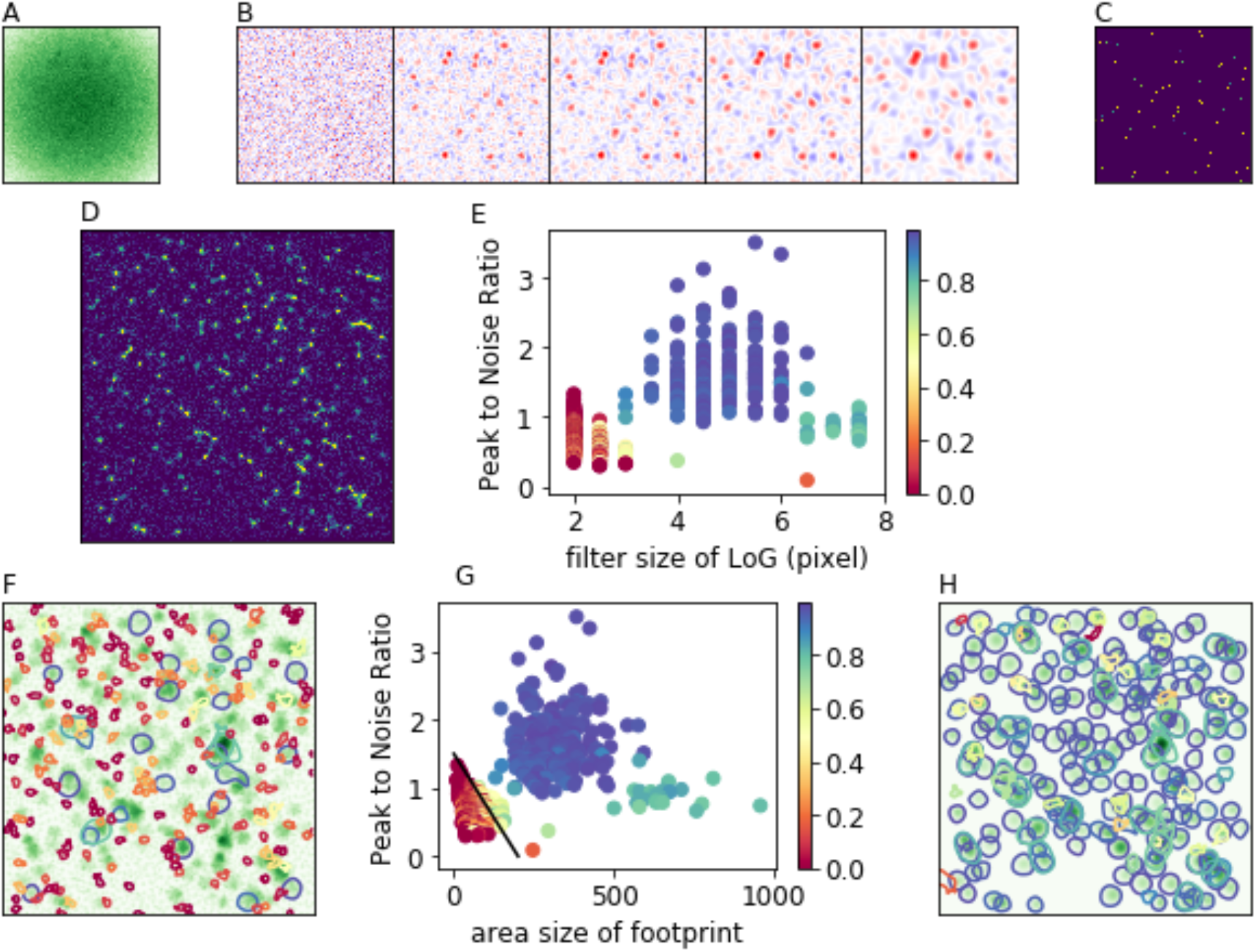
Detection of cell candidates. (A) Example frame of data. (B) Filtered images of (A) using the Laplacian of Gaussian filter with 2-, 4-, 6-, 8-, and 10-pixel scales. (C) Peaks of (B) in three dimensions (position and scale). (D) Peaks in all frames. (E) The scale and strength of peaks are distributed into two groups, and the color illustrates the cosine similarity to the grand truth cell footprint. (F) Randomly selected footprints corresponding to peaks in (E). (G) Area size and peak strength of the footprints in (F). (H) Selected footprints by the line indicated in (G).

Next, peaks representing neighborhoods are detected from all detected peaks in all frames (Fig. 2D). The selected peaks are derived from artifacts due to noise as well those as from real cells, and their distribution differs largely depending on the scale of the LoG and the signal strength (Fig. 2E). Then, we applied the watershed method to each selected peak to segment out the footprint (Fig. 3F). Footprints can be also roughly divided into noise and cells based on the peak value and area size (Fig. 2G).

**Figure 3.**
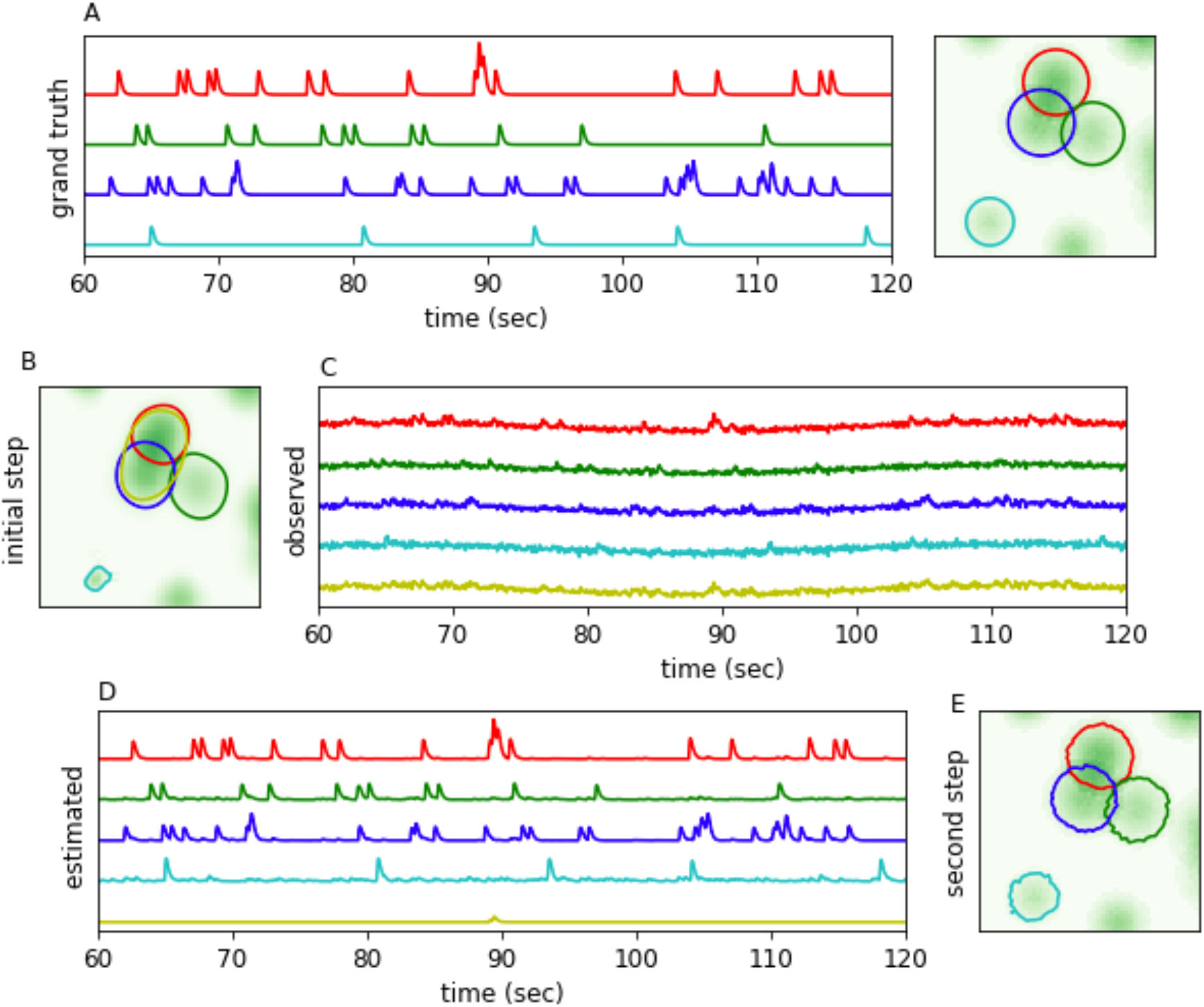
Spatial and temporal components are iteratively improved by enhanced constrained nonnegative matrix factorization. (A) Grand truth of four cells. (B) Initial footprints. (C) Observed activities averaged over initial footprint. (D) Estimated calcium activities from observed activities with the information of overlap. (E) Reconstructed footprints. Even a footprint with a weak correlation (cyan) can be reconstructed using the temporal and spatial steps. Contrarily, false positives (i.e., yellow) are automatically removed.

Using simple criteria (as shown in Fig. 2G), all 200 cells can be detected, suppressing the number of duplicated footprints (Fig. 2H). In this step, LoG processing can be performed independently for each frame as well as each peak to remove peripheral regions; thus, the calculation can be effectively performed via parallelization.

### Enhanced CNMF improves iterative temporal and spatial components with automatic relevance determination

The calcium time series averaged over a footprint can be easily calculated (Fig. 3A and B), and the cell activities can be estimated from the average time series and data on overlapped footprints (Fig. 3C) using the CNMF framework. In this step, we introduced a method that can estimate calcium signal simultaneously with whole baseline, which was conventionally done separately. A priori knowledge of spike-calcium dynamics as impulse responses and the firing rate of each cell as a parameter are also modeled similarly as the original CNMF (see Algorithm for Temporal Steps in Method Details).

Conversely, in the spatial steps, the footprints can be estimated from the estimated calcium activities and a priori knowledge of cell shape and size (Fig. 3D). By applying the time and space steps in order, the footprint is obviously closer to the grand truth (Fig. 3E). Conversely, the estimated calcium signal for the false-positive footprint has a relatively small amplitude, and it is automatically removed during the space step.

### Shrink estimation of the baseline enhances the robustness and stability of the automatic relevance determination

As described in the previous section, it is expected that by repeating the time and space steps, the cell shape and activity estimations will be accurate. Indeed, it is possible to reproduce nearly all 200 cells in the artificial data by removing false positives via iteration using the proposed method (Fig. 4A and B). In the proposed method, even if the spike firing frequency and sparsity parameter for cell shape are changed, almost equivalent results are obtained (Fig. 4C).

**Figure 4.**
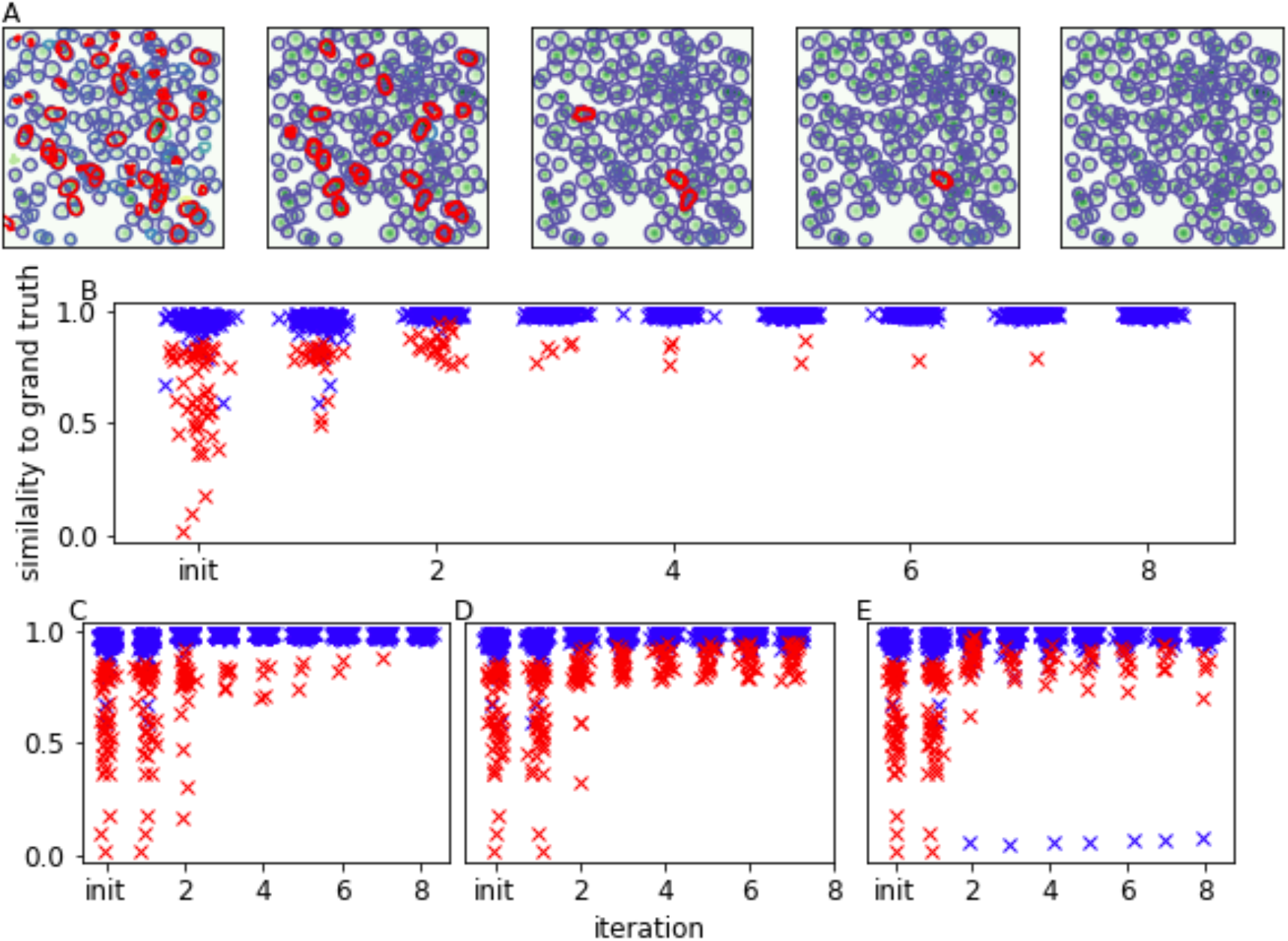
Convergence in multiple iterative steps. (A) False positives were eliminated by repeating the iteration (initial state, 2-, 4-, 6- and 8-th iteration, respectively). (B) Correlation to the grand truth. (C) Results of different parameter settings. (D–E) Results without shrink estimation corresponding to (B) and (C). Without shrink estimation of the baseline, false positives were not eliminated, and correlation with the grand truth was decreased.

**Figure 5.**
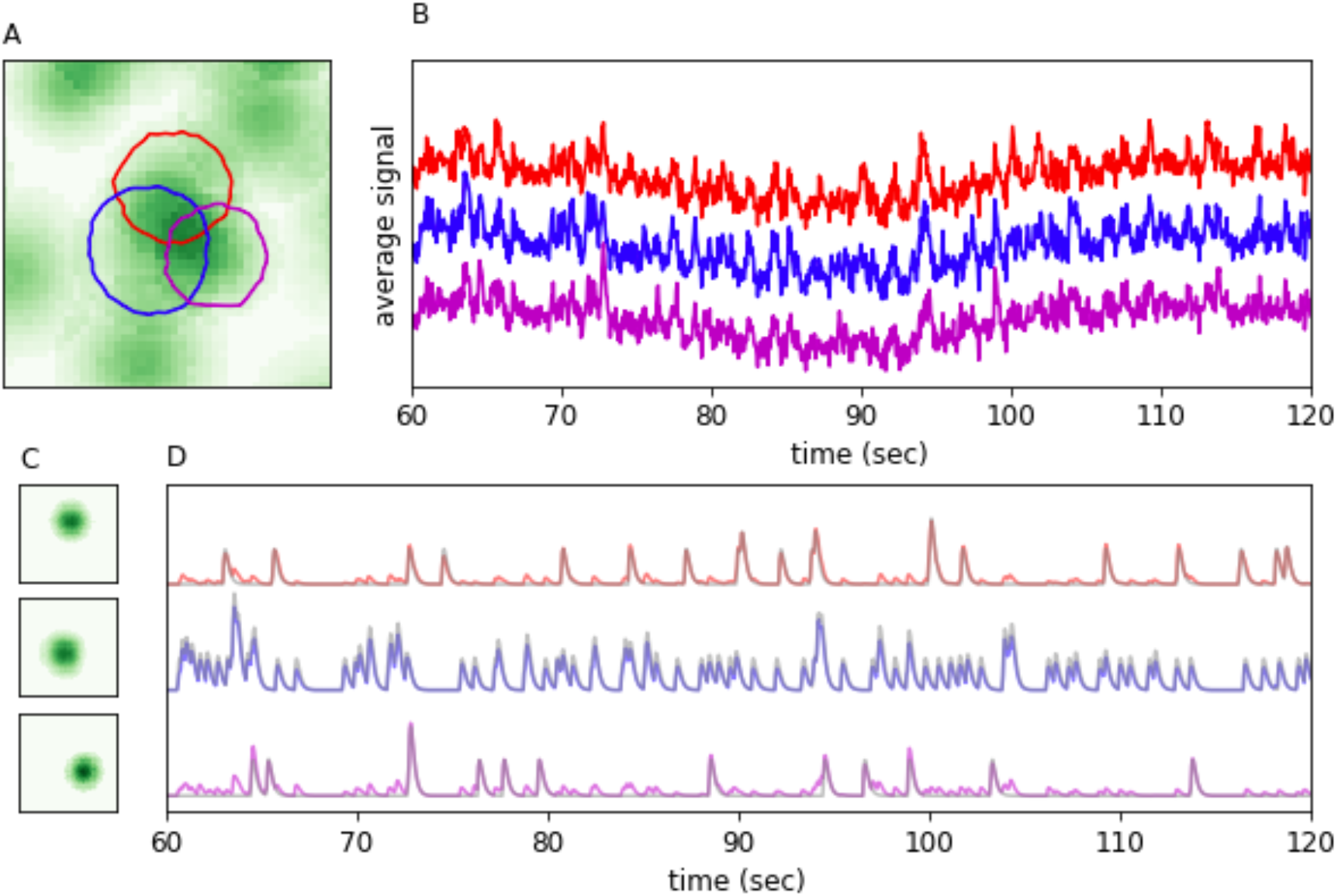
Accuracy of the estimation of calcium activities. The activities to three overlapped cells can be reconstructed by removing crosstalk from the observed signals. The peak-to-noise ratio of three cells are 1.32, 1.18 and 1.36, respectively.

Conversely, when shrink estimation of the baseline is not performed, false positives are not properly eliminated (Fig. 4D), and for some parameters, correlation with the grand truth deteriorates due to repetition, resulting in divergence of the calculation (Fig. 4E). This stabilization property is obtained by balancing sparse parameters and the baseline reduction estimation while simultaneously estimating cellular elements and the baseline. In addition, the number of conditions of the matrix appearing in the iterative method for solving the optimization problem at each step improves, and thus, the number of calculations is reduced.

As a result, calcium time series can be estimated with high accuracy via repetition. Even when particularly strong overlap exists, it is possible to stably improve and converge the cell shape and activity time series.

### Application of the method to hippocampal CA1 data

It is necessary to confirm the properties for the artificial data using real data. Thus, the proposed method was applied to recorded hippocampal data. First, to detect cell candidates using the LoG filter, comparison with ROI data confirmed via visual recognition revealed results similar to the artificial data regarding footprint size and signal intensity, and footprint matching with the manual recognition result appeared together in the upper right (Fig. 6A). Using this distribution, cell candidates were selected as the initial conditions, and repeating the time and spatial steps covered all ROIs via visual recognition, although cell candidates that could not be confirmed via visual recognition remained (Fig. 6B).

**Figure 6.**
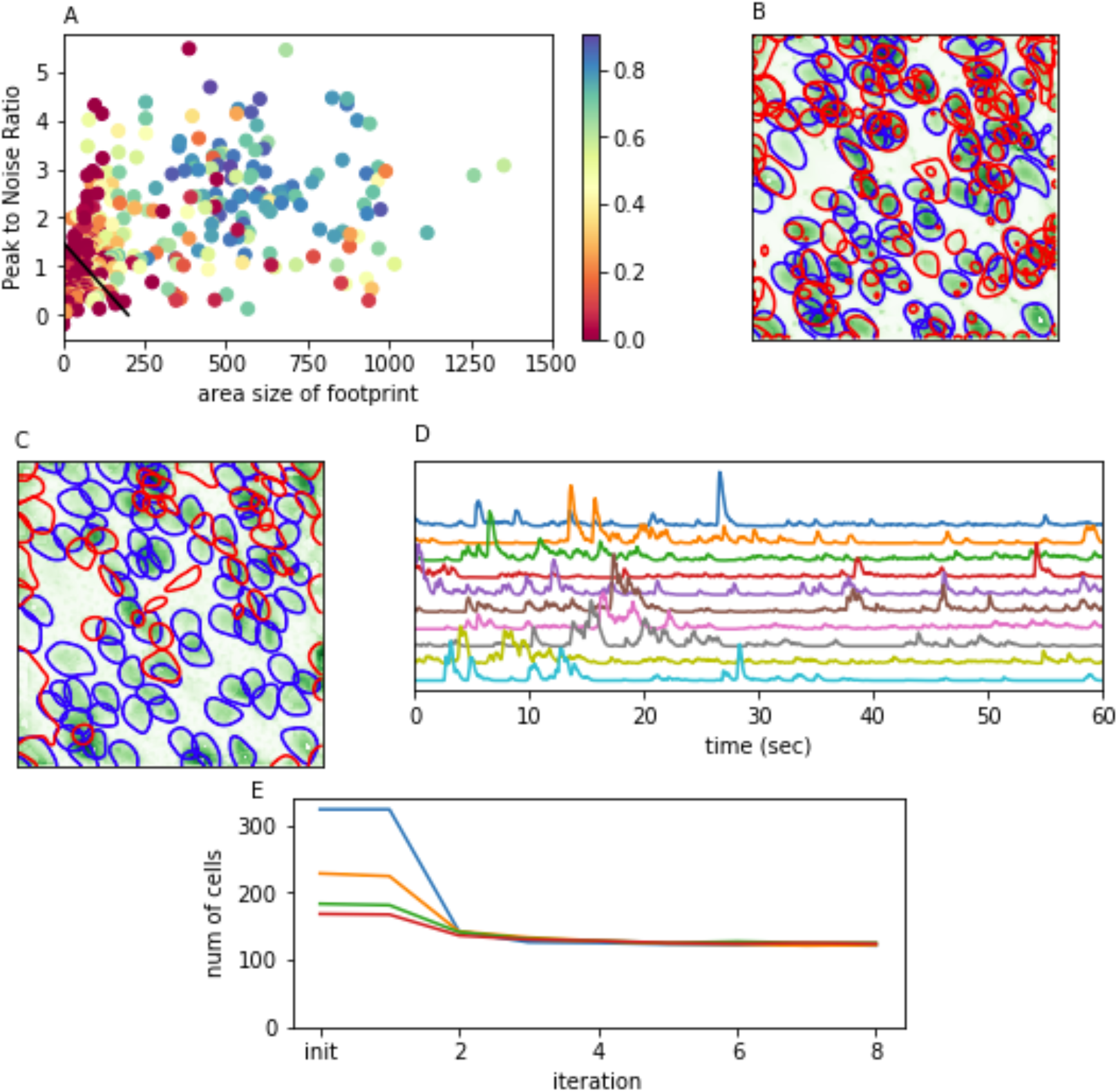
Result for hippocampus data. Peak and footprint detection using the Laplacian of Gaussian filter. (A) Distribution of cell sizes and peak signal intensities. (B) Selected cell candidates. (C-D) Footprint and time series obtained as the final result are presented. (E) The calculation was repeated while changing the initial state.

By applying the iterative algorithm from the obtained initial condition, results including 18 cell candidates in addition to 84 manually obtained cells were obtained (Fig. 6C). In addition, almost the same number of cells was detected when more initial candidates were selected or even when the sparseness parameter was changed (Fig. 6D).

## DISCUSSION

In this paper, we proposed a footprint initialization method and baseline shrinkage estimation for the CNMF framework and its efficient calculation method. The latter method stabilizes the convergence when repeating time and space steps and confirms that the robustness of the result is obtained. The proposed method is considered effective for several other methods already proposed in the CNMF framework. Conversely, it is also expected that further improvements in accuracy can be obtained by introducing a local background. In our current implementation, FISTA was used as a solver of the sparse model, but it is expected that higher speed can be achieved using alternating direction method of multipliers (ADMM) or linearized ADMM, which can make good use of the relationship between the spike and calcium. Simultaneous estimation of prior parameters can be performed using LARS, but it is not necessarily effective in terms of reducing the calculation time because of the relatively large number of nonzero elements.

## ACKNOWLEDGMENTS

This work was supported by the Core Research for Evolutional Science and Technology (CREST) program (JPMJCR13W1) of the Japan Science and Technology Agency (JST), KAKENHI (23220009) from the JSPS, Grant-in-Aid for Scientific Research on Innovative Areas “Memory dynamism” (25115002) from the MEXT, the Mitsubishi Foundation, the Uehara Memorial Foundation, the Takeda Science Foundation support to K.I. T.F. is supported by Brain/Minds project from the MEXT, KAKENHI (15H04265) from the JSPS, Grant-in-Aid for Scientific Research on Innovative Areas “Memory dynamism” (16H01289) and “Artificial Intelligence and Brain Science” (17H06036) from MEXT. Y.H. is supported by RIKEN, Human Frontier Science Programme, a Grant-in-Aid for Scientific Research on Innovative Areas “Memory dynamism” (16H01292) and “Synapse Pathology” (22110006) from the MEXT. T.T. is supported by KAKENHI (26870577) from the JSPS.

## EXPERIMENTAL MODEL AND SUBJECT DETAILS

### Hippocampal CA1 data

All procedures involving the use of animals complied with the guidelines of the National Institutes of Health, and they were approved by the Animal Care and Use Committee of the University of Toyama and the Institutional Committee for the Care and RIKEN Animal Experiments Committee and Genetic Recombinant Experiment Safety Committee. Thy1::G-CaMP7-T2A-DsRed2 mice that express the fluorescent calcium indicator protein G-CaMP7 and the red fluorescent protein DsRed2 under the control of the neuron-specific Thy1 promoter were used. Details on the generation and characterization of the transgenic mice will be described elsewhere (Sato et al., submitted). The mice were maintained on a 12-h/12-h light-dark cycle (lights on 7:00 am) at 24 ± 3°C and 55 ± 5% humidity, provided *ad libitum* access to food and water, and housed with littermates until 1–5 days before surgery.

We performed hippocampal surgery for gradient refractive index (GRIN) relay lens setting as previously described (Barretto, et al., 2011; Ghosh, et al., 2011; Ziv, et al., 2013). All surgeries were conducted on approximately 12-week-old male Thy1::G-CaMP7-p2A-DsRed mice on a C57BL/6J background. Mice were anesthetized with a pentobarbital solution (80 mg/kg of body weight; intraperitoneal injection), and the fully anesthetized mice were placed in a stereotactic apparatus (Narishige, Japan). To set the cannula lens sleeve (outer diameter, 1.8 mm; length, 3.6 mm; Inscopix, CA), craniotomy was performed with a diameter of 2.0 mm. The cylindrical column of the neocortex and corpus callosum above the alveus covering the dorsal hippocampus was aspirated using a 27-gauge blunt drawing up needle with saline. The cannula lens sleeve was gently placed on the alveus and fixed to the edge of the burr hole with bone wax, which was melted using a low-temperature cautery. The cannula lens sleeve targeted the right hemisphere (AP 2.0 mm, ML 1.5 mm at center). After setting the anchor screws onto the skull, we covered the skull with dental cement, which fixed the cannula lens sleeve to the skull and anchor screws.

Approximately 3–4 weeks after surgery, mice were anesthetized with isoflurane (1.5–2%), and a GRIN lens (outer diameter, 1.0 mm; length, 4.0 mm; Inscopix) was inserted into the cannula lens sleeve and fixed with ultraviolet-curing adhesive (Norland, NOA 81). The integrated microscope (nVista HD, Inscopix) (Ghosh, et al., 2011) with a microscope baseplate (Inscopix) was placed above the GRIN lens, at which G-CaMP7 fluorescence was observed. The microscope baseplate was fixed with the head of the anchor screw using dental cement, via which the GRIN lens was shaded, after which the integrated microscope was detached from the baseplate. The GRIN lens was covered by attaching the microscope baseplate cover to the base plate until calcium imaging was performed.

Calcium imaging was performed during the light cycle, and calcium events were captured at 20 Hz using nVista acquisition software (Inscopix). The integrated microscope was re-attached to the baseplate, and then mice were introduced into a novel context consisting of a cylindrical chamber (diameter × height: 180 × 230 mm^2^) with a white acrylic floor and walls covered with black tape. The captured movie was processed as previously described (Kitamura, et al., 2015). After motion correction using Mosaic software (Inscopix) and division of each image on a pixel-by-pixel basis using a low-passed [r = 20 pixels] filtered version on ImageJ, the *dF/F* signal movie was prepared using Mosaic software.

To manually detect calcium signals in each cell, ROIs for cell locations were selected using Mosaic software.

## METHOD DETAILS

### Finding cell candidates from movie data using LoG filters

The recoded movie data can be represented by the matrix 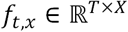, where *T* is the number of frames and *X* is the number of pixels in the target area. To detect cell-like blobs in each frame, we applied a scale-normalized LoG filter with multiple scales and searched local peaks larger than threshold 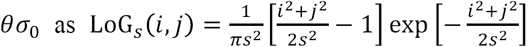,

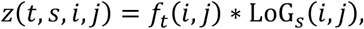

where the position is denoted as *x* = (*i,j*), *θ* is the S/N ratio, and *σ*_0_ is the noise level estimated using the following equation:

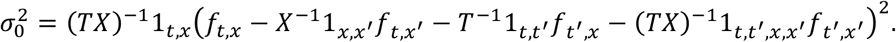

Note that the representation of the sum in the same index is omitted by the notation of the tensor.

Then, duplicated peaks were removed from all detected peaks, and the footprints were segmented around the peak. Finally, we can choose cell candidates according to the footprint size and peak intensity using the following procedure.

1 For each *t*

1-1 Find local peaks{(*t*, *s*, *i*, *j*)|(*s*, *i*, *j*) is local maximum of *z*(*t*, *s*, *i*, *j*), *z*(*t*, *s*, *i*, *j*) > *θσ*_0_}

2 Remove duplicated peaks

2 — 1 Sort detected peaks (*t*,s*, i*,j\**) by signal strength *z*(*t*\*,*s*\*, *i*,j\**)

2-2 Initialize *mask*(*i, j*) ← 0

2 — 3 Pop (*t*, i*,j*,s\**), to step 3 if empty

2 — 4 If *mask*(*i, j*) = 0, then select and *mask*(*i, j*) ← 1 for neighbor (*i, j*) of (*i*, j\**)

3 Obtain the footprint from the peak (*t*, s*, i*,j\**) using a watershed-like algorithm

3-1-l Push (*i*, j\**) to the list

3-1-2 Pop (*i, j*)with the largest (*t*, s*, i, j*) while list is not empty

3-1-3 Push neighbor pixels (*i′, j′*) of (*i, j*) if 0 < *z*(*i′,j′*) *≤ z*(*i, j*)

### Mathematical model for calcium imaging data

We assumed that the imaging field contained a total number of *K* neurons. The cell activity of the *k*-th neuron is the direct product of the nonnegative “spatial footprint” *a*_*k,x*_ and “temporal fluorescence intensity” *v*_*k,t*_ for this neuron. The movie signal *f*_*t,x*_ is observed as the sum of the baseline *b*_*t,x*_ and the cell activities *a*_*k,x*_*v*_*k,t*_ with noise *ε*_*t,x*_ as follows:

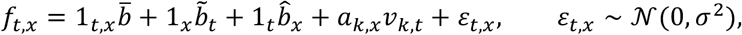

where *ε*_*t,x*_ obeyed the normal distribution with variance *σ*^2^.

We assumed that the baseline signal level of the calcium imaging data could be separated into constant 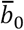, temporal 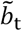, and spatial 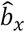 components as follows:

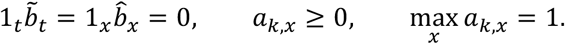

The temporal component *v*_*k,t*_ is the impulse response *g*_*t,τ*_ of the nonnegative spiking activity *u*_*k,τ*_. The calcium sensitivity to the spike signal is denoted by *s*_*k*_.

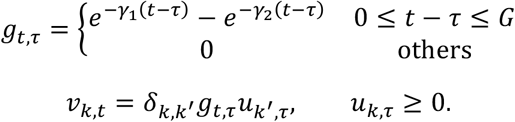

### Baseline prior

We assumed a normal distribution as a prior distribution of the baseline for both time and space. The variance of the prior distribution is set to a constant multiple of the error variance as follows:

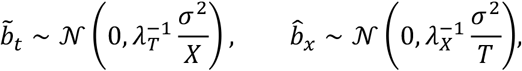

At this time, the log posterior distribution becomes

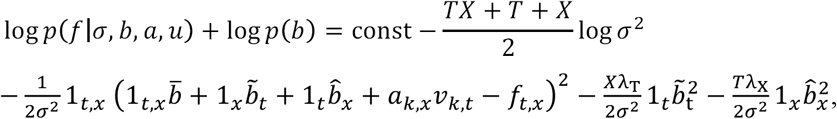

and minimizing this by *b\** and *σ\** results in the following:

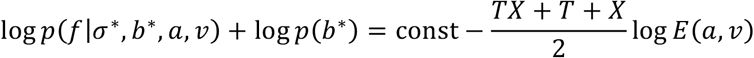

where

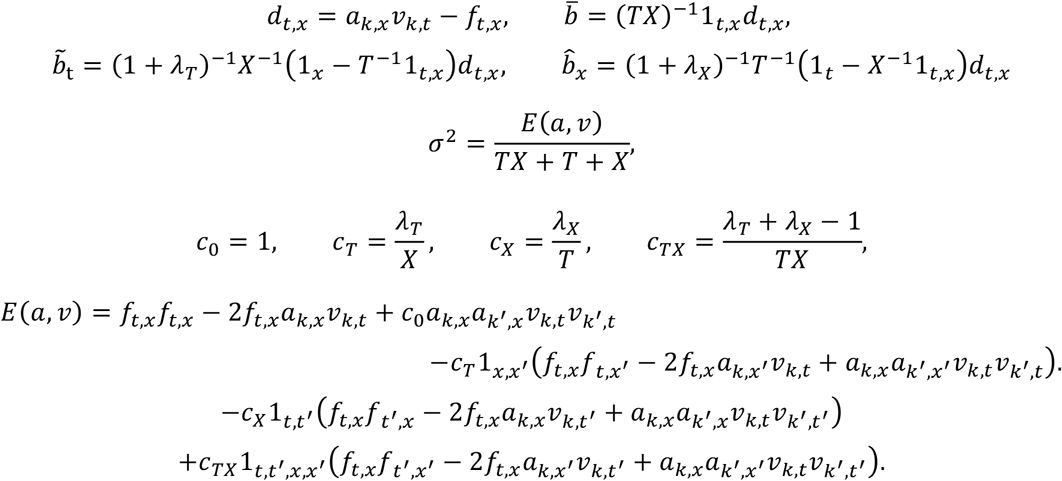

### Main solving problem

Our main solving problem can be simply defined as

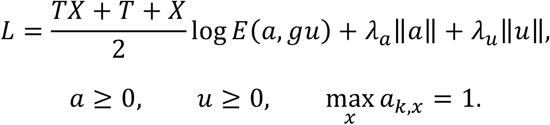

The main problem can be solved via iterative temporal and spatial steps. We used an accelerated proximal gradient method called FISTA (Beck & Teboulle, 2009) for each step.

### Algorithms for temporal steps

We used a normalized norm for the penalty term *λ*_*u*_|*u*|| as

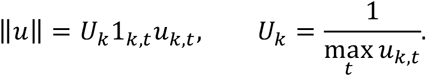

If we fixed *a\**, the main solving problem could be simplified as

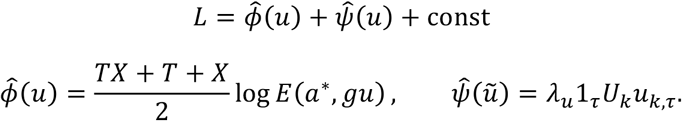

The target function *L*(*u*) can be decreased via the update step of the proximal gradient method as

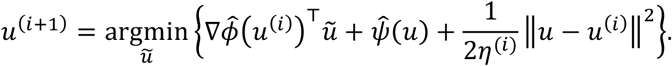

The right term can be divided into the element-wise function

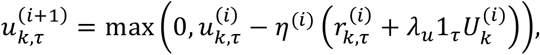

where

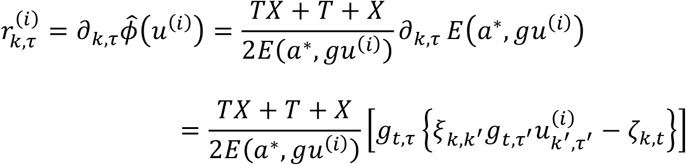

and

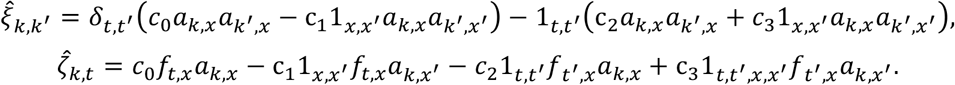

### Algorithms for the spatial steps

If we fix *u\**, the main solving problem can be simplified as

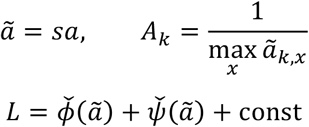

The target function *L*(*ã*) can be decreased via the update step of the proximal gradient method as

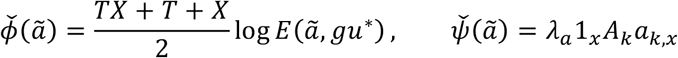

The right term can be divided into an element-wise function as follows:

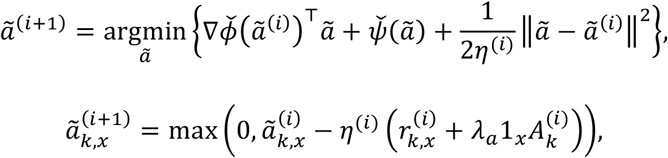

where

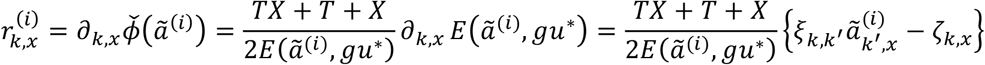

and

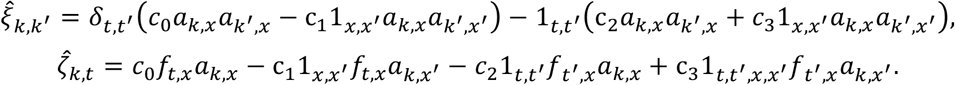

### Segment footprints

Despite not introducing a spatial structure in the optimized model, the result obtained in the space step is suitable as the footprint of the cell. Nonetheless, increased restriction of the cell area makes the iterative convergence more stable. In this study, the cell area was limited using the same LoG filter and segment method as described for the initialization. However, in the initialization, the output was also set to the value of the LoG filter. In this case, the result of the calculation of the region using the LoG filter was used as the image mask for the originally estimated footprint.

### Remove false positives and merge existing components

It is possible to aggressively eliminate cell candidates using the difference of the score. When the score is increased by removing cell candidate *k*, we can remove *k* as follows:

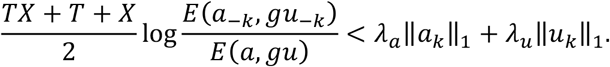

We can also merge similar components. The merged component of *l* and *m* can be obtained via simple NMF as follows:

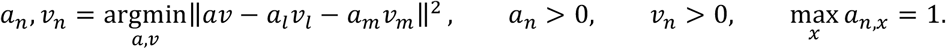

We can replace the cell candidate pair *l*, *m* to the merged components *n* when the score increases as follows:

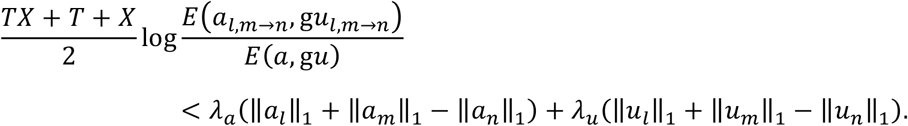

### Overall framework

1. Initial footprints
2. Temporal step
3. Spatial step
4. Segment footprints
5. Temporal step
6. Remove small candidates (optional)
7. Merge similar candidates (optional)
8. Return to 3 if not converged

At this point, we will explain the process used to numerically solve the aforementioned temporal and spatial steps. These technical issues are not essential for the accuracy of the obtained solutions, but they are crucial for efficiently solving the time-consuming maximization problem for a large data set.

Solving the primary duality problem is a standard method for solving a maximization problem in the presence of constraints. In this method, we switch the roles of the primary objective functions and the constraints, noting that the maximization with respect to the model parameters and the minimization with respect to Lagrange multipliers should occur simultaneously at the optimal point.

### Parameter settings

We roughly approximate the prior parameter of the temporal component *λ*_*u*_ using ratio between the sampling *S* and typical firing rate *R* as follows:

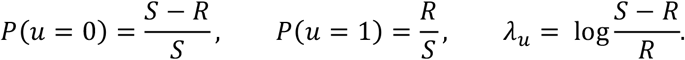

The prior parameter of the spatial components was approximated using the ratio between the total number of pixels *X* and typical footprint size *A* as follows:

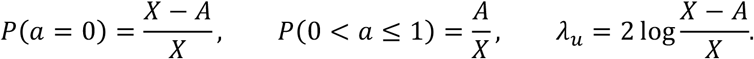

### Complexity analysis

In the spatial step, the time cost is mainly determined by two components, namely *O*(*KTX*) for calculating *f*_*t,x*_*a*_*k,x*_ and *O*(*IK*^2^*X*) for iteratively calculating 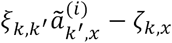. The number of iterations is always smaller than 20 for both simulated and real data.

In the temporal step, the update procedure includes an additional step to translate from u to v via the impulse response of length G. Therefore, the time cost is *O*(*KTX + IK*^2^*T + IKGT*). The number of iterations is always smaller than 200 for both simulated and real data.

### Implementation

All analyses were performed with custom-written Python code using numpy/scipy (van der Walt, et al., 2011) and scikit-image (van der Walt, et al., 2014) packages. The implementation uses Dask (Dask Development Team, 2016; Rocklin, 2015), MKL (Intel), and TBB (Intel) for parallelized effective computation.

### Simulation data

In the simulation data for Fig. 1 to Fig. 5, we simulated 200 cells in 12,000 frames of 300 × 300 pixels. Firing rates of cells followed a log-normal distribution (*μ_k_ ~ LN*(0.5,0.4)) and the number of spikes were between 24-283. The peak-to-noise ratio also followed a log-normal distribution (*s*_*k*_ ~ *LN* 1,0.2)) and were between 0.7-2.1 as a result. The footprints were two dimensional Gaussian function of random center and uniform distributed standard deviation (4-6 pixel). The spatial and temporal baseline were quadratic and sinusoidal function, respectively and the simulated data can be described as 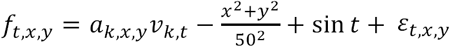.

